# The Implications of Over-Estimating Gene Tree Discordance on a Rapid-Radiation Species Tree (Blattodea: Blaberidae)

**DOI:** 10.1101/717660

**Authors:** Dominic A. Evangelista, Michael A. Gilchrist, Frédéric Legendre, Brian O’Meara

**Affiliations:** Department of Ecology and Evolutionary Biology, The University of Tennessee, Dabney Hall, 1416 Circle Dr., Knoxville, TN 37996, USA; Institut de Systématique, Evolution, Biodiversité (ISYEB), Muséum national d’Histoire naturelle, CNRS, Sorbonne Université, EPHE, UA, 57 rue Cuvier, CP50, 75005 Paris, France; National Institute for Mathematical and Biological Synthesis, Knoxville, TN, 37996 USA

**Keywords:** Congruence, incomplete lineage sorting, tree error, mutation selection, substitution model, evolutionary model, Insecta

## Abstract

Patterns of discordance between gene trees and the species trees they reside in are crucial to the debate over the superiority of coalescent or concatenation approaches to tree inference. However, errors in estimating gene tree topologies obfuscate the issue by making gene trees appear erroneously discordant with the species tree. We thus test the prevalence of discordance between gene trees and their species tree using an empirical dataset for a clade with a rapid radiation (Blaberidae). We find that one model of codon evolution (FMutSel0) prefers gene trees that are less discordant, while another (SelAC) shows no such preference. We compare the species trees resulting from the selected sets of gene trees on the basis of internal consistency, predictive ability, and congruence with independent data. The species tree resulting from gene trees those chosen by FMutSel0, a set with low discordance, is the most robust and biologically plausible. Thus, we conclude that the results from FMutSel0 are better supported: simple models (i.e., GTR and ECM) infer trees with erroneously high levels of gene tree discordance. Furthermore, the amount of discordance in the set of gene trees has a large effect on the downstream phylogeny. Thus, decreasing gene tree error by lessening erroneous discordance can result in higher quality species trees. These results allow us to support relationships among blaberid cockroaches that were previously in flux as they now demonstrate molecular and morphological congruence.

## Introduction

Debates over the relative merits of concatenation or coalescent tree inference are ongoing (Gatesy and Springer 2014; Roch and Steel 2015; Edwards, et al. 2016). Edwards, et al. (2016) posit that, the debate addresses the differences among gene tree topologies - concatenation represents a “special case” of coalescence where all gene tree topologies are presumed to be the same. However, concatenation and coalescence usually yield largely similar topologies (Leache and Rannala 2011; Edwards, et al. 2016) but see (Mendes and Hahn 2018). This suggests the “special case”, or nearly identical cases (i.e., where the vast majority of gene trees are the same), are not especially rare for many nodes on a phylogeny. We investigate how common gene tree to species tree (GTST) discordance is in under different evolutionary models. We approach this using an empirical dataset of Blaberidae cockroaches, a phylogeny that is difficult to infer due to a, presumably, rapid radiation (Legendre, et al. 2017).

The biological processes contributing to GTST discordance [i.e., incomplete lineage sorting (ILS), hybridization and lateral gene transfer (Huang, et al. 2017; Thawornwattana, et al. 2018)] can be accounted for through coalescent theory (e.g., Maddison 1997). In many studies, ILS is presumed to be responsible for all discordance between gene trees and the species tree (e.g., Chou, et al. 2015), since hybridization and lateral gene transfer events are deemed more rare (Copetti, et al. 2017). Yet, gene tree inaccuracy is also expected to contribute to GTST discordance. On reason being that modelling mutational variance within a gene tree is difficult (Huang, et al. 2010). Some approaches account for uncertainty in gene trees when inferring a phylogeny through the coalescent process (Rannala and Yang 2017; Flouri, et al. 2018; Wang and Nakhleh 2018) but most methods do not (Kubatko, et al. 2009; Than and Nakhleh 2009; Liu, et al. 2010; Leache and Rannala 2011). In either case, improved accuracy in gene trees would likely lead to more accurate species trees since coalescent methods are highly sensitive to small changes in gene trees (Shen, et al. 2017; Wang and Nakhleh 2018) or rogue gene trees (Reid, et al. 2014).

In this study, it is assumed that if estimated gene trees contain some phylogenetic error those errors will usually contribute to increased discordance with the species tree. With this assumption in mind, we test the abundance of GTST discordance using three evolutionary models to minimize gene tree error. Two selection-based codon models (SelAC; Beaulieu, et al. 2019; FMutSel0; Yang and Nielsen 2008) and one nucleotide model (GTR; Tavaré and Miura 1986) are compared.

Since genes have limited character information, they may be greatly affected by small errors from unrealistic modelling of nucleotide evolution. Codon models are inherently more complex than nucleotide models, to reflect the complexity of natural processes acting on evolution of protein coding genes. Thus, codon models are thought to improve phylogenetic inference (e.g., Goldman and Yang 1994; Wang, et al. 2014; Arenas 2015; Doud, et al. 2015; Sealfon, et al. 2015). Modelling codon data is also thought to be more effective than modelling amino-acid evolution (Arenas 2015). SelAC and FMutSel0 are similar in that they include selection to model mutational evolutionary history. However, FMutSel uses a single substitution matrix and combines effects of both stabilizing and frequency dependent selection [via their use of omega to model the non-synonymous to synonymous substitution rate dN/dS; see Beaulieu, et al. (2019)]. In contrast, SelAC solely models stabilizing selection, but assumes the strength of stabilizing selection scales with gene expression (Drummond and Wilke 2008)and that substitution rates depend on the physio-chemical distance between the optimal amino acid for a site and its alternatives. As a result, rather than using a single substitution matrix, SelAC employs 20 different matrix families, but whose elements are parameterized using a small number of parameters.

Improving our ability to identify discordance of gene trees in species trees improves the tractability of a classic phylogenetic problem: rapid radiations (Giarla and Esselstyn 2015). The problem stems from both low phylogenetic signal from the short period of the radiation and the increased effect of ILS on short branches (Giarla and Esselstyn 2015; Mendes and Hahn 2018). Our approach (Fig. 1) addresses this issue from both sides. First, by using two codon models (SelAC and FMutSel0) that, in theory, are more biologically plausible than simpler models (e.g., GTR, ECM). Second, by using a coalescent method (Mirarab and Warnow 2015). Coalescent methods account for ILS, which can confound resolution on short internodes when it is not accounted for.

**Figure 1.**
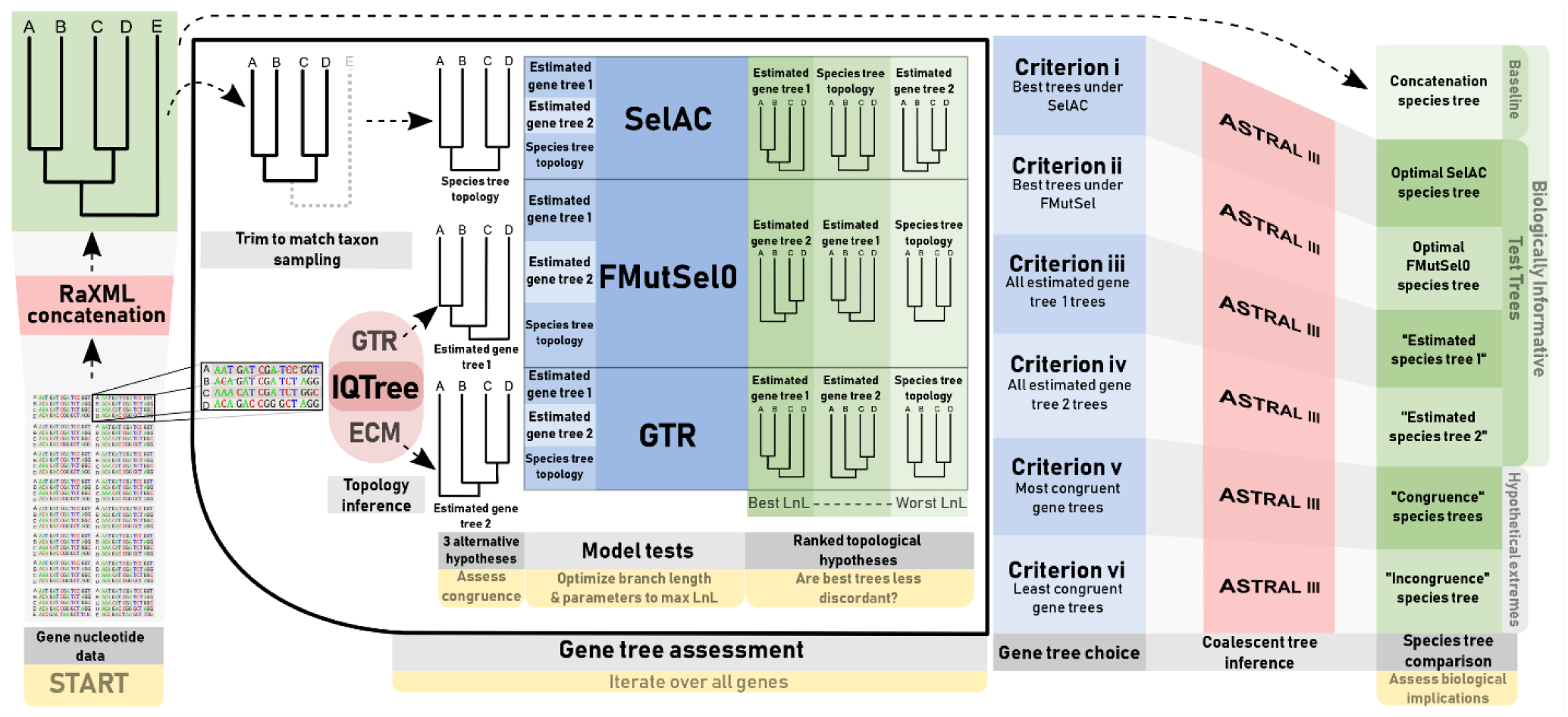
A schematic diagram of the workflow for testing the abundance of gene tree discordance under three models, and downstream affects in the species tree.

We test this using challenging biological data from the cockroach family Blaberidae, whose major lineages diversified 100 Ma in a period of only 30-40 Myr. (Legendre, et al. 2015; Evangelista, et al. 2019). Evangelista, et al. (2017) illustrated that most subfamilies in Blaberidae have not been placed consistently. Yet, some of the consistent relationships have not been supported by more recent evidence (Legendre, et al. 2017; Evangelista, et al. 2018; Evangelista, et al. 2019 unpublished data). Some specific examples are as follows. Which lineage is recovered as sister to the remaining Blaberidae has been inconsistent and highly dependent on taxon sampling (Evangelista, et al. 2018). The origins of Panchlorinae and Neotropical Epilamprinae as being ancient or more recent is under contention (Legendre, et al. 2017; Bourguignon, et al. 2018; Evangelista, et al. 2018; Evangelista, et al. 2019 unpublished data). Also, taxa with many morphologically derived traits like *Diploptera punctata, Pycnoscelus* spp., and *Aptera fusca* also have no consistent placement across phylogenies (Legendre, et al. 2017, Evangelista, et al. 2019; Evangelista, et al. 2018; Evangelista, et al. 2019 unpublished data). Some groups [e.g., Paranauphoetinae, Panesthiinae and Perisphaerinae in Anisyutkin (2003) or Blaberinae and Zetoborinae in Grandcolas (1993)] have morphological hypotheses for their relationships that have not been strongly supported by molecular data (Legendre, et al. 2017; Bourguignon, et al. 2018; Evangelista, et al. 2018; Evangelista, et al. 2019 unpublished data). Sometimes, inconsistent topologies have prevented interpretation of interesting phenotypic diversity (e.g., Madagascar hissing cockroaches; Inward, et al. 2007; Legendre, et al. 2017; Evangelista, et al. 2018; Evangelista, et al. 2019 unpublished data). While most previous studies have had poor sampling with regard to Blaberidae, Legendre, et al. (2017) and to a lesser extent Evangelista, et al. (2019 unpublished data) have greatly improved this. Yet, they still recover largely incongruent topologies.

## Materials & Methods

All genetic data were taken from Evangelista, et al. (2019 unpublished data). From their 265 protein coding loci a subset of 100 loci were chosen based on minimizing locus rate heterogeneity (calculated in that study). Each alignment was further reduced to an ingroup sampling of 42 Blaberidae species and 14 other Dictyopteran outgroups to allow for maximal data completeness.

### Species Trees and Gene Trees

A baseline concatenation tree was inferred using all taxa and all 265 loci from Evangelista, et al. (2019 unpublished data) using their exact procedure. Briefly, GTR and GTR+G were applied in a partitioned analysis of a concatenated alignment in RaXML {Stamatakis, 2014 #2431} with a single topology assumed for all partitions. Partitioning scheme and model choice were optimized in PartitionFinder 2 {Lanfear, 2016 #2721}. This topology was subsequently used as an approximate species-tree topology (“concatenation topology”/”concat.”). While it is not a perfect topology, it is reliable as a standard because it uses an abundance of data. Compared to our further analyses below, the concatenation topology was inferred using nearly ten times more nucleotide data, seven more ingroup taxa, and almost three times more taxa overall.

The tips of the concatenation topology were trimmed (Paradis, et al. 2018) to match the taxon sampling of the gene trees and species trees. Two gene-trees were then inferred from each locus in IQ-TREE (Nguyen, et al. 2015). The first was inferred using the GTR model (“estimated gene tree 1”/ “Est.GT.1”), a median-approximated gamma distribution with 4 rate categories (G4), optimized nucleotide frequencies (FO), and the thorough “allnni” search option. The second tree was inferred with the codon model ECMS05 (Schneider, et al. 2005) (“estimated gene tree 2”/ “Est.GT.2”), G4, and a nucleotide frequency model determined by IQ-TREE’s built-in model finder (Kalyaanamoorthy, et al. 2017) under the AICc (Sugiura 1976) optimality criterion (see supplementary material S1.1 for further details on model choice). We consider all Est.GT.1 or all Est.GT.2 trees as being in the same “model family” since they share substitution models. Rate heterogeneity was also modelled using the gamma distribution (four rate categories) but with a mean-approximation method. All three topologies were identically rooted with an outgroup taxon (*Metallyticus splendidus*, a solumblattodean, or *Ectoneura* sp. depending upon taxon sampling of that gene).

The Robinson-Foulds (RF; Robinson and Foulds 1981) and the path distance (Kuhner and Yamato 2015) among all topologies (Concat., Est.GT.1, and Est.GT.2) for each locus were measured using the function treedist in the R package Phangorn (Schliep 2011). RF distance is the more intuitive metric as it gives a measure of the number of clades shared between trees. However, when even one rogue taxon is present it can give very high distances, which is undesirable. The Path distance metric is much less sensitive to rogue taxa but is a less biologically meaningful descriptor of tree differences {Kuhner, 2015 #2710}. The path distance is the sum of differences in minimal species pair path lengths between trees {Kuhner, 2015 #2710}. All distances were greater than 0, indicating that all three hypothetical phylogenies for each locus were different. Yet, since RF and path distances compare different aspects of tree shape, a given tree can be the more discordant tree under one metric but the less discordant tree under the other. We discarded all loci for which this was the case (∼34% of cases).

### Model Tests

Preliminary tests (supplementary material S2.2) showed that FMutSel0 preferred trees of one model family (i.e. model family bias; Table S.2.2.1). To account for this bias we limited our tests to 40 loci, where the least discordant trees were equally distributed among both model families. These loci had an average of 44 species per alignment (min. 34, max. 50).

Branch lengths for each of the three trees for each locus were optimized again using the SelAC software package (Beaulieu, J.M., O’Meara, B.C., et al. 2019). We fit: GTR+G4+FO, SelAC+GTR+G4+FO+amino acid optimization (AAO), and MutSel+GTR+G4+FO+AAO. In both cases, estimated branch lengths from IQ-TREE were used as starting values and optimization chain was run for four sets of initial conditions. The first set used six iterations of 1000 evaluations each, and the subsequent three sets used only four iterations of 1300 evaluations each. Other parameters used were max.tol.edges = 1.4 and tol.step = 2.3, parallelized over two processors. The highest log-likelihood (lnL) of the four independent optimizations was chosen as the preferred tree. The number of independent optimizations and the chain length were deemed effective because test runs showed that difference in log-likelihood between the independent runs was an order of magnitude less than the average difference in lnL between treatments (trees).

If the topology with the best lnL under a given model (i.e., SelAC, FMutSel0, GTR) was also the estimated tree with the lowest discordance, as determined by both RF and path distances, the trial was assigned a 1, otherwise a 0. Z-tests were used assess if the models identified the least discordant tree more often than expected from randomness and if the number of trees chosen were random with respect to the model family of the tree (i.e., Est.GT.1/GTR or Est.GT.2/ECM). A test was also done to see if lnL’s were skewed for trees of a certain model family or for more discordant trees. For the former, the lnL of Est.GT.1 was subtracted from that of Est.GT.2 and compared the median of the distribution to the null expectation of a randomized sample of delta lnL’s with a Kolmogorov-Smirnoff test. The same was done for the difference between the lnL of the least discordant estimated gene tree and the most discordant. Finally, we examined if the difference in lnL’s (ΔlnL) between gene tree 1 and gene tree 2 was dependent on the magnitude of hypothesis distinctness as measured by either tree-distance between the estimated gene trees or the difference in topological distance to the concatenation tree. This was done with a linear regression. All type-I error thresholds were predefined at 0.05 and calculations were done in Mathematica 10 (Wolfram Research 2016).

### Testing the Implications on the Species Tree

The downstream implications of the choice of gene tree topologies were tested through a coalescent tree inference in ASTRAL-III (Mirarab and Warnow 2015). ASTRAL-III is an appropriate software because it is consistent with the coalescent process, it allows gene trees to have different taxon sampling, and does not co-estimate the gene tree and species tree. Four test trees were inferred from the following sets of gene trees: all estimated gene tree 1 topologies (Est.ST.1 tree), all estimated gene tree 2 topologies (Est.ST.2 tree), all trees chosen as optimal by SelAC (SelAC tree), and all chosen as optimal by FMutSel0 (FMutSel0 tree). These were also compared to the phylogeny inferred from concatenation of: all loci (concatenation tree; the baseline), the most discordant trees (max. discord tree) and the least discordant ones (min. discord tree).

A “distance score” was calculated to categorize the species trees as more consistent with maximal or minimal discord in the gene tree sample. Distance score was calculated as (*D*_*max*_ − *D*_*concat*_) + (*D*_*max*_ − *D*_*minDisc*_) + (*D*_*maxDisc*_) where D_max_ is the maximum distance between any species tree pair, D_concat_ is the distance to the concatenation tree, D_minDisc_ is the distance to the minimum discordance tree, and D_maxDisc_ is the distance to the maximum discordance tree. The distances used were RF and path distances rescaled between zero and one and then added together. High distance scores indicate that a tree is more consistent with minimally discordant gene trees. Then topological comparisons were made to determine which recovered relationships were attributable to maximal or minimal discord.

The plausibility of the tested species trees tells which gene tree sample yields the most realistic evolutionary scenario. Thus, each species tree was evaluated under three criteria: internal consistency, predictive power, and congruence. High internal consistency was defined as high mean local posterior probability and high mean clade frequency in the underlying gene trees. Predictive power was assessed by counting how often relationships in the species tree were found in gene trees (estimated under both models as described above) for 60 loci not used in the coalescent tree inferences. These were compared statistically by randomly resampling the 60 gene trees with replacement for 100 trials. At each trial the mean of the frequency of occurrences were taken each time and the distribution of means were compared via a Z-Test as implemented in the R package BSDA (Arnholt 2019). Congruence was defined as presence of predefined control nodes (see supplementary material S1.2), presence of previously hypothesized relationships, and presence of morphological support for relationships. See supplementary material S2.3 for justification and evidence for each. Also, see table S2.3.1 for the methodology and results for a test of plausibility given concatenated datasets (Shimodaira 2002).

## Results

The 40 alignments had a median parsimony score of 1076.5 (min. 260, max. 4711; based on the Est.GT.1 topologies). Gene tree to species tree (GTST) discordance was ubiquitous because no estimated gene trees came within 20 RF from the concatenation tree (Fig. 2). This level of discordance is approximately equivalent to two random clade swaps in a 45 taxon tree. Each estimated topology was unique; they were never less than 20 RF distance from each other (Fig. 2). The GTR model tended to estimate slightly less discordant trees compared to the ECM models (78% of cases; Fig. 2 a, c, e). With few exceptions, the gene trees were more similar to each other than they were to the concatenation tree (Fig. 2 b, d, f).

**Figure 2.**
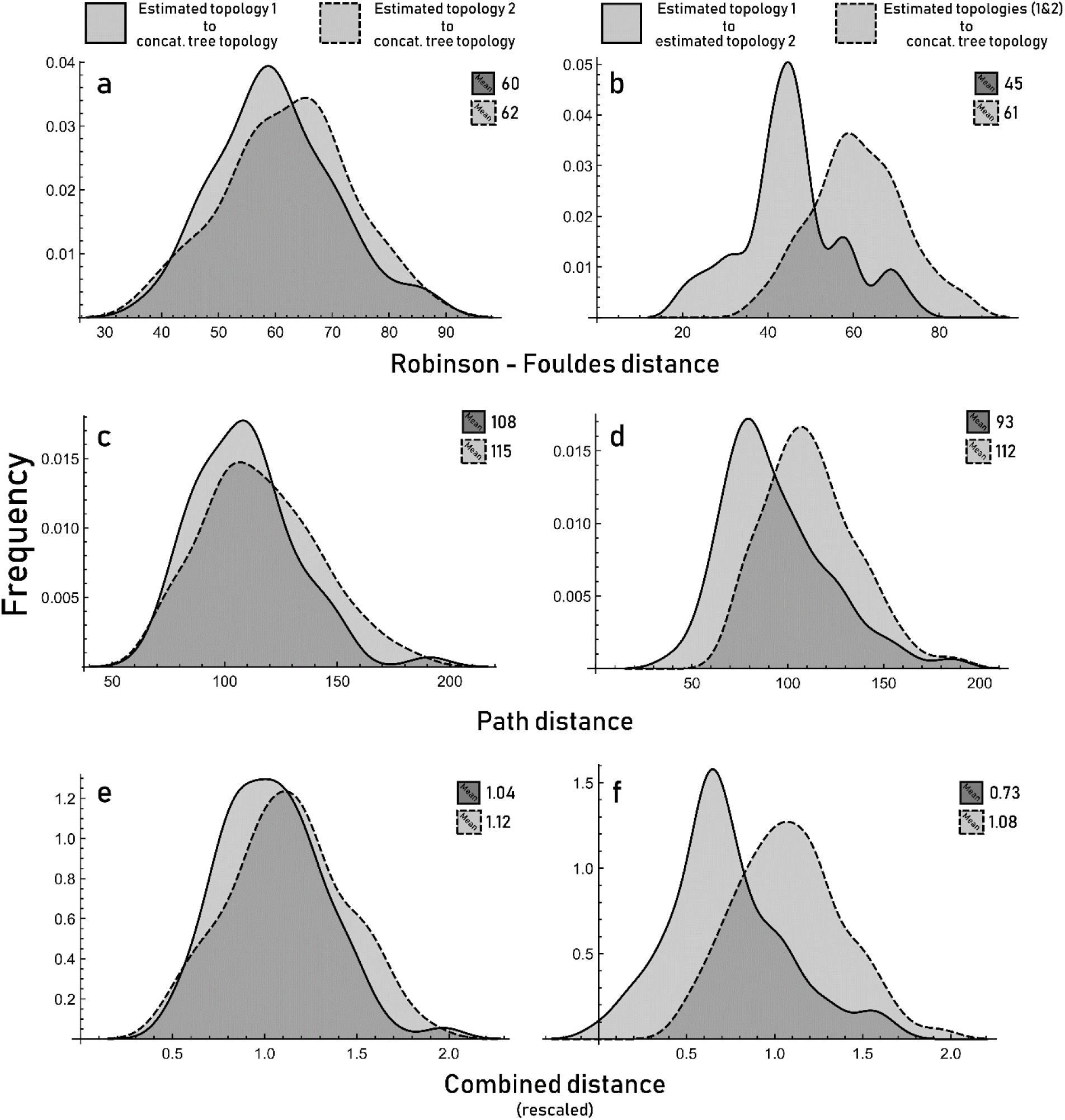
Histograms showing the distribution of topological distances among possible gene tree topologies. Robinson-Foulds distances are given in (a, b), path distance in (c, d) and a combined and rescaled distance is given in (e, f). The distances from the two estimated gene trees to the concatenation tree topology (all with equal taxon sampling) are shown in the left panels (a, c, e); the distance between both estimated gene trees and the mean distance to the concatenation tree topology are shown in the right panels (b, d, f). Est.GT.1 tended to be slightly less discordant than Est.GT.2. Estimated gene trees were more similar to each other than they were to the concatenation tree.

### Model Tests

Only FMutSel0 had a statistically identifiable preference for less discordant trees (Fig. 3). GTR and SelAC both assigned the best likelihood (lnL) to the least discordant tree 60% of the time, which was not more than expected by random choice (p>0.05). FMutSel0 was significantly more likely to choose the least discordant tree (p = 0.006) and chose them 70% of the time. Both GTR and FMutSel0 preferred gene trees estimated under the GTR model (i.e., Est.GT.1) 65-70% of the time (p<0.05). SelAC showed no such model family bias.

**Figure 3.**
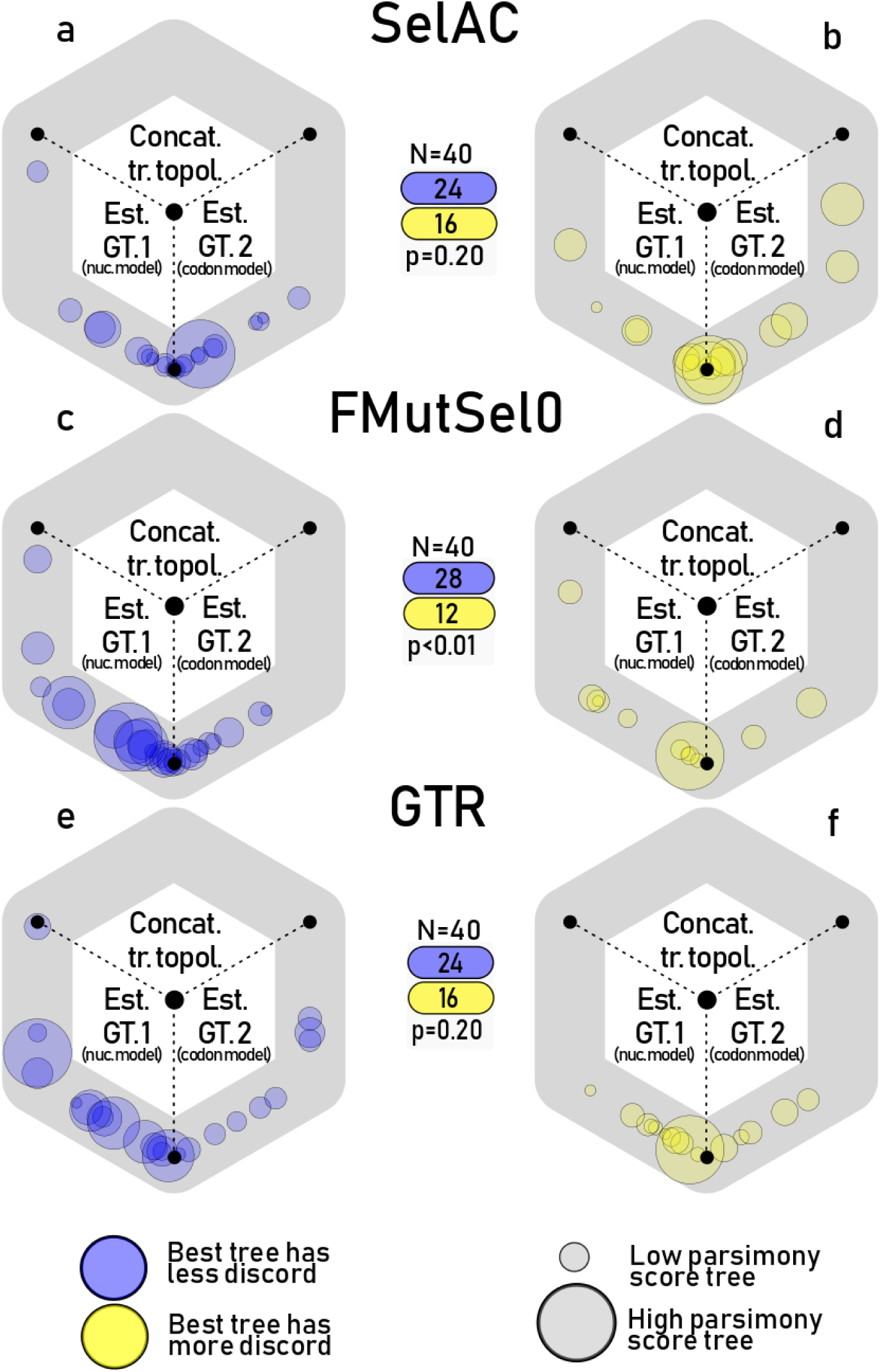
Results from the tests of 40 genes under three models: SelAC (a, b), FMutSel0 (c, d), and GTR (e, f). Circles along the hexagon each represent a genomic locus, its position represents the relative likelihood of the three possible tree topologies. For instance, a circle in the bottom left (8 o’clock) position represents a locus where estimated gene tree 1 was much more likely than the other two topologies. A circle just to the left of the bottom position, (6-7 o’clock) represents a slight preference for the same tree relative to that of estimated gene tree 2. Circle radii indicate the parsimony scores of loci. The results are separated by cases where the models favored the least discordant (blue) or most discordant (yellow) trees. For nearly all tests, the concatenation topology was the least favored, indicating that some discordance was always preferred. Only FMutSel0 demonstrated a non-zero preference for the least discordant gene tree.

The rank order of likelihoods from the model tests inform on the most likely gene trees. The difference in the magnitude of the lnLs give further insight into the behavior of the model tests. Examining these differences shows that GTR and FMutSel0 are still biased by the model family of the tree but FMutSel0 did not have better lnLs for less discordant trees (Fig. 4). Thus, while FMutSel0 preferentially chose trees with lower discordance, the lnL magnitudes resulting in those choices were not significantly different from those expected under random choice.

**Figure 4.**
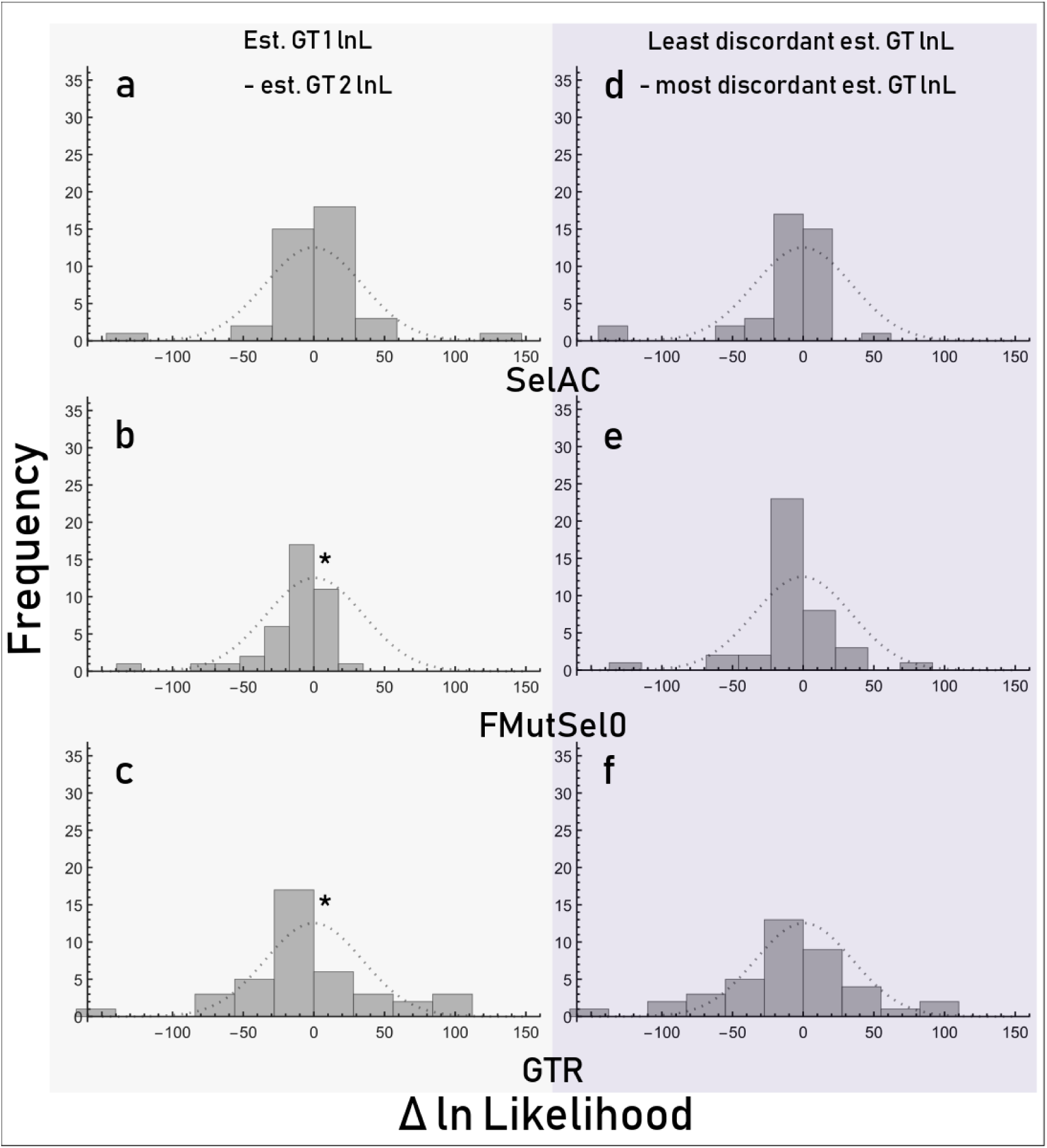
Histograms showing the distribution of delta ln-likelihoods for the indicated trees under the three models. Dashed lines represent the null distribution expected as determined based on a randomized resampling of all the data. * indicates statistically significant deviation from the mean of the null expectation as determined from a Kolmogorov-Smirnoff test (α = 0.05). This test only considers the magnitude of preference for a certain estimated gene tree. The magnitudes of lnL’s from FMutSel0 and GTR were slightly biased towards Est.GT.1.

Finally, there was no relationship between how distinct the estimated gene trees were from each other (Fig. 5) and the lnL they received. The magnitude of discordance to the concatenation tree had no relationship to the magnitude of lnLs from any of the three models (Fig. 5 d-f). With SelAC, the difference in lnL did show a relationship to the topological distance between the two estimated topologies (Fig. 5 a). However, when the two outliers on the left are removed, the trend entirely disappears (p=0.91, R^2^<0.01, m=-0.02).

**Figure 5.**
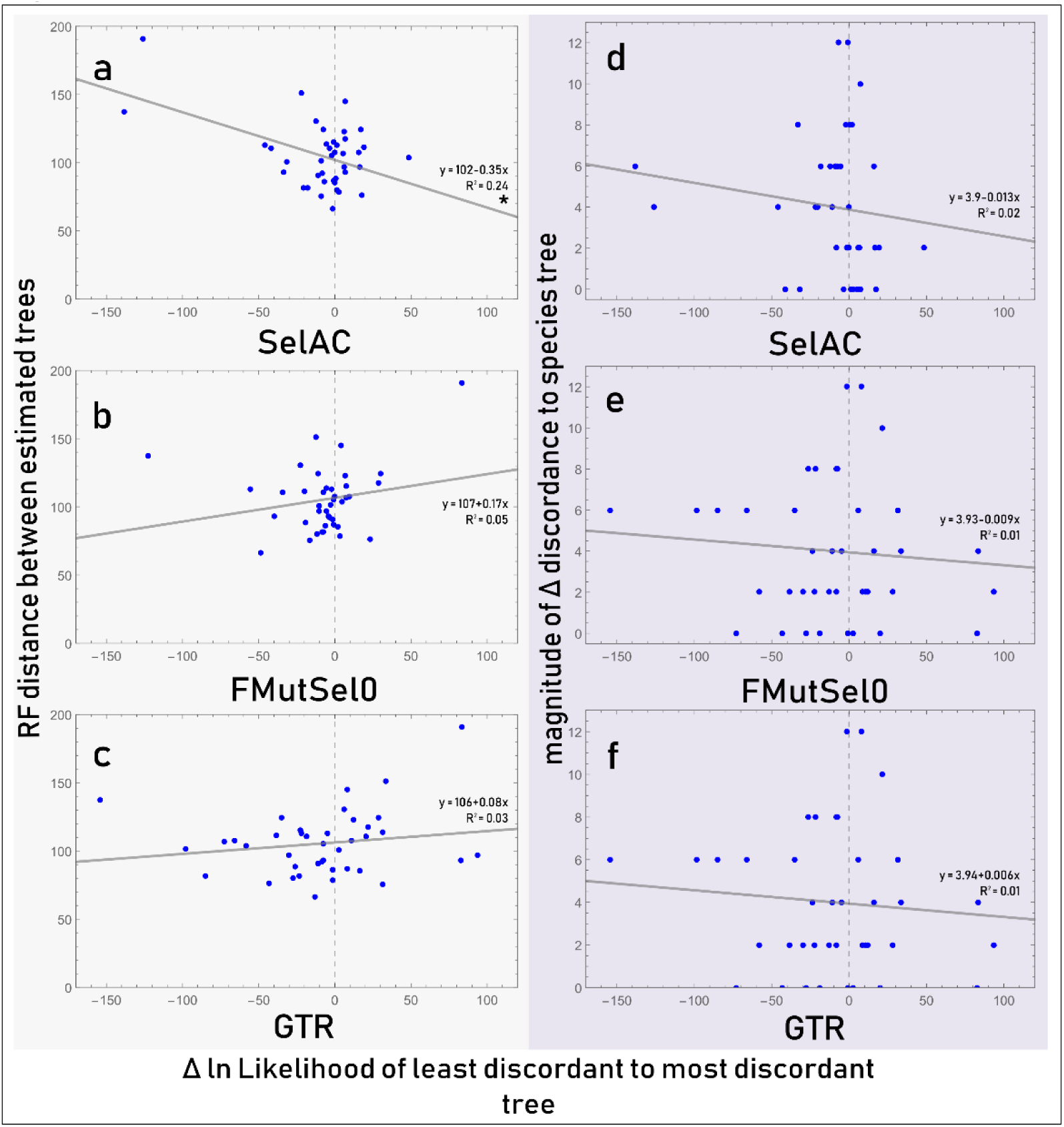
Relationship between preferences for the least discordant estimated gene tree and the amount of difference between the estimated gene trees. Left panels show the differences between the two estimated gene trees (a, b, c) and right panels show the magnitude of difference in discordance among the two estimated gene trees. Data is fit with a linear model and statistical significance determined at an α level of 0.05. The only statistically significant regression (a) was found to be entirely driven by the presence the two outlying points on the left.

### Effects of GTST Discordance on Species Trees

Specifics of the species tree topologies, support for relationships under different scenarios, predictive ability of the species trees, and detailed comparison of their overall plausibility are given in supplementary material S2.2 and S2.3. Figure S2.2.1 shows the differences in the topologies of the four test trees (Est.ST.1, Est.ST.2, SelAC, FMutSel0), a best-case scenario (min. discord), worst case scenario (max. discord) and baseline tree (concatenation).

Relationships in the FMutSel0 species tree were the most consistent with relationships found in the more congruent gene trees. The FMutSel0 species tree had the best “distance score” (1.22) due to its high similarity to the baseline and min. discordance trees and low similarity to the max. discordance tree (Table 1). Among the four test trees, the SelAC tree had the poorest “distance score” (0.76) due to its high similarity with the max. discord tree and low similarity to the min. discord and baseline (concatenation) trees (Table 1).

**Table 1.** Comparison of species trees.

| Species Tree | Data | Gene tree model | Analysis | Distance score<sup>1</sup> | Selected relationships score<sup>2</sup> |
| --- | --- | --- | --- | --- | --- |
| Concatenation | 265 loci | NA | RaXML<br>(partitioned,<br>GTR) | 2.88 | 20 |
| Min.<br>discordance | 40 least discordant gene<br>trees | GTR or<br>ECMS05 | ASTRAL | 1.86 | 13 |
| Max.<br>discordance | 40 most discordant gene<br>trees | GTR or<br>ECMS05 | ASTRAL | 0.33 | -9 |
| Est.ST. 1 | 40 Est.GT.1 topologies | GTR | ASTRAL | 1.17 | 1 |
| Est.ST. 2 | 40 Est.GT.2 topologies | ECMS05 | ASTRAL | 0.85 | -2 |
| SelAC | 40 SelAC chosen gene<br>trees | GTR or<br>ECMS05 | ASTRAL | 0.76 | 0 |
| FMutSel0 | 40 FMutSel0 chosen<br>gene trees | GTR or<br>ECMS05 | ASTRAL | 1.22 | 2 |

*Note:* The FMutSel0 species tree is most consistent with the more congruent baseline tree and the concatenation topology. Est.ST.2 is the least consistent.
<sup>1</sup> Distance score is a composite of scaled RF and path distances from the baseline trees (concatenation, min. discordance, max. discordance). The value is higher when distance from the concatenation tree and min. discord tree is lower and distance to the max. discord tree is higher.
<sup>2</sup> Selected relationship score is higher when a tree shares a certain relationship (see supplemental data) with the concatenation or min. discordance tree and is lower when it shares a relationship with the max. discordance tree.

To determine test tree plausibility, we examined: internal consistency, power to predict independent gene tree topologies, and congruence with previous studies. The species trees with the highest internal consistency (mean local posterior probability, clade frequency in constituent gene trees) were: Est.ST.1 (0.79, 36.8%), FMutSel0 (0.77, 36.7%), SelAC (0.77, 36.6%), and Est.ST.2 (0.76, 35.5%) (Table 2). The species trees that predicted the highest number of relationships found in an independent sample of 120 gene trees (from 60 loci) were: Est.ST.2 (29.1% ± 2.2%), FMutSel0 (28.9% ± 1.9%), Est.ST.1 (28.8% ± 2.0%), and SelAC (28.7% ± 2.0%) (Table 2). Est.ST.2 and FMutSel0 predicted relationships in gene trees an indistinguishable rate (p > 0.05) but each did so more frequently than the Est.ST.1 and SelAC species trees (p < 0.05).

**Table 2.**
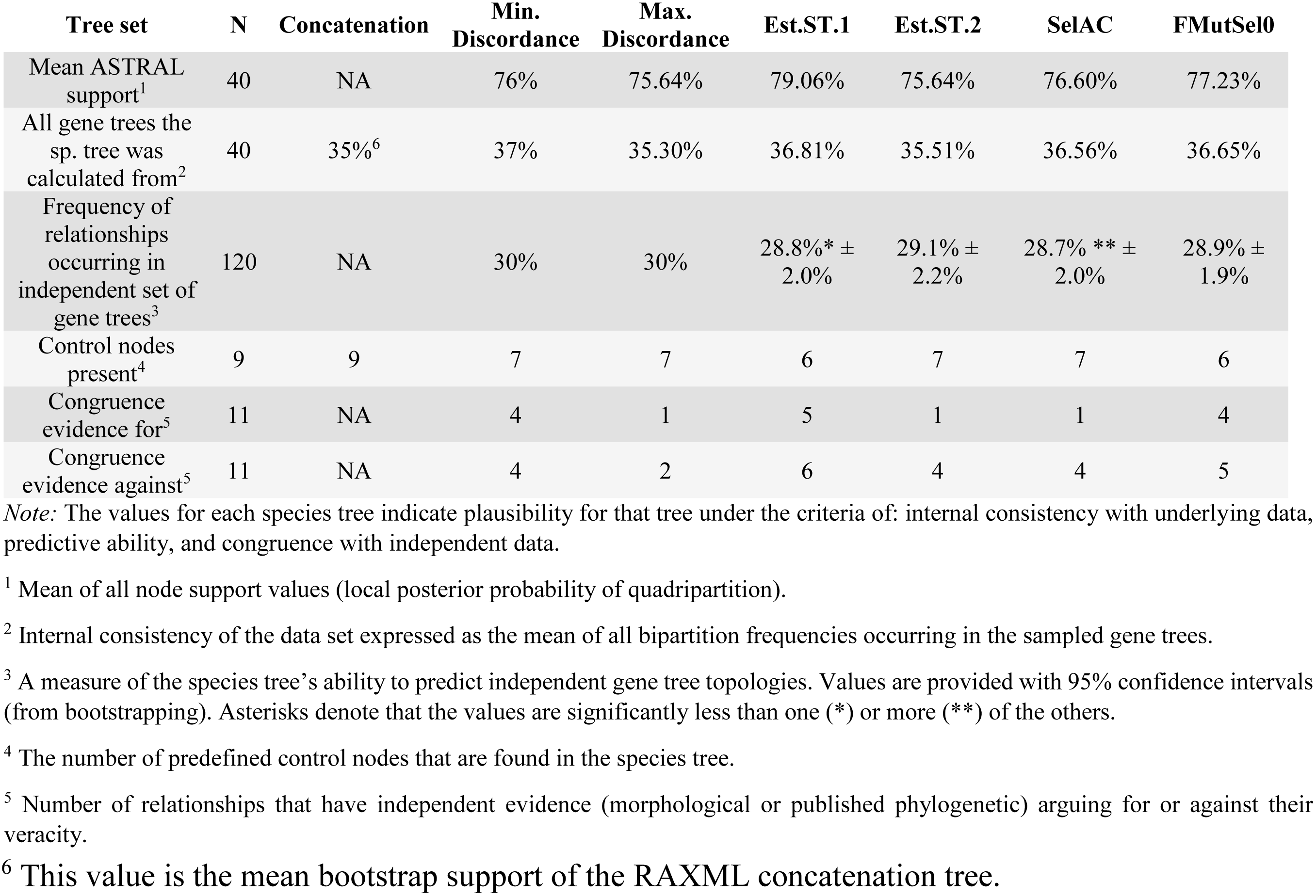
Metrics to evaluate the plausibility of each species tree.

As one test of congruence, we looked for a set of control nodes - relationships known with high certainty (see supplementary material S1.2). Only the concatenation tree had all nine control nodes present. Each test species tree had six (Est.ST.1, FMutSel0) or seven (Est.ST.2, SelAC) control nodes present - differing with each other only in which lineage was sister to Blattellidae *sensu* Evangelista, et al. (2019 unpublished data).

As a second test of congruence, we evaluated which relationships in the trees had prior support from published morphological studies or independent phylogenetic analyses. Of the 11 Blaberidae taxa whose sister relationships were examined (Fig. S2.3.3), the relationships recovered in the Est.ST.1 and FMutSel0 trees were most plausible (Table 2; see supplementary material S2.3 for specific information). In Est.ST.1 there were five relationships with putative independent support and six relationships with independent data conflicting with them. In the FMutSel0 tree, there were four relationships with supporting data and five relationships with conflicting data. The Est.ST.2 and SelAC trees each had one relationship supported and four relationships conflicted.

## Discussion

The results from the FMutSel0 model show that our hypothesis is correct with respect to GTST discordance (Fig. 3). However, FMutSel0 did not prefer less discordance in all trees (Fig. 4 e) but preferred less discordant trees more often and more discordant ones (Fig. 3 c). SelAC did not select less discordant trees at a statistically significant rate in either way (Fig. 3 a 4 d). One possible confounding factor is that FMutSel0 and GTR were both biased towards selecting estimated gene tree 1 (Fig. 3, Table S2.1.1) even when discordance was balanced between the two sets of gene trees.

GTR was expected to favor estimated gene tree 1 in most cases since this tree was inferred with GTR in IQ-TREE. However, this is the opposite of what was expected for FMutSel0, which would be predicted to have a bias towards estimated gene tree 2, which was inferred under the codon model. One explanation for this is a trade-off between shared model traits and biological realism. In theory, SelAC should be less susceptible to the effects of model family bias because it accounts for nucleotide, amino-acid, and protein changes simultaneously. This would explain the observed patterns of bias except that FMutSel0 showed a bias away from the codon model topology (Est.GT.2). An explanation for this reverse bias is unclear but could be due to poor performance of the empirical codon model (ECM; Schneider, et al. 2005; Kosiol, et al. 2007) when only given single genes (Delport, et al. 2010). Since the results show no indication that FMutSel0 or SelAC are choosing gene trees based on shared model characteristics, we assume their choices are more related to plausibility of the gene trees under the model given the data (i.e., biological realism under the assumptions of the model).

Coalescent inference using gene trees selected by FMutSel0 yield a species tree topology that was the most plausible considering: consistency with its own data (quadripartition support; Table 2), ability to predict independent gene tree topologies (Table 2, Table S2.3.2), and congruence with independent data (Table 2; Table S2.3.3). FMutSel0, however, validated one control node less than two other species trees (Table 2). Since the results overwhelmingly favor the FMutSel0 tree, we favor those results with respect to GTST discordance and with respect to the species tree.

Thus, we conclude that lessening GTST discordance results in a stronger species tree and likely reduced gene tree error. In the debate between coalescence and concatenation, this result supports a middle ground but does favor coalescence. Trees were never lacking discord altogether (with respect to any species tree and any estimated gene tree). Yet, GTST discord was usually overestimated. Thus Edwards, et al. (2016)’s “special case”, where all gene trees have the same topology for most nodes in a phylogeny, is indeed rare but discord is still less abundant than simple models suggest.

Our results show that error is abundant in estimated gene trees. Since individual genes may not have enough character information to inform accurate gene trees (Roch, et al. in press), model-derived gene tree errors doubly compound this problem. This emphasizes the importance of utilizing approaches that co-estimate gene trees and species trees to account for errors (e.g., Höhna, et al. 2016; Rannala and Yang 2017; Flouri, et al. 2018; Wang and Nakhleh 2018). Indeed, these errors have a meaningful effect on the downstream species tree if they are unaccounted for (Table 1, Fig. S2.2.1, S2.3.3). Of course, joint estimation of species trees and gene trees is also vulnerable to errors from ineffective modelling of alignments (Reid, et al. 2014). Given this, studies whose results hinge largely on the prevalence of ILS may need to be examined more closely (Copetti, et al. 2017).

Finally, our protocol suggests that extracting signal from short loci may be more effective with a mutation selection codon model since the FMutSel0 tree was higher quality than the others. This is somewhat surprising considering the biological shortcomings of this model. While both FMutSel0 and SelAC ignore the effects of selection on codon usage, this fact is of particular concern for FMutSel0 since it includes a dN/dS term in its formulation. The use of dN/dS to model sequence evolution in this way has been criticized since it only effectively describes scenarios with positive or negative frequency dependent selection (Hughes, et al. 1990; Kryazhimskiy and Plotkin 2008; Wolf, et al. 2009; Mugal, et al. 2014; Beaulieu, et al. 2019), but it is a simpler model that is easier to fit. MutSel models of codon evolution also don’t effectively model sites under positive selection, which leads to erroneous estimates of omega (Spielman and Wilke 2015).

Est.ST.1 also performed well (Table 2) and its topology was consistent with the species tree inferred from all the least discordant trees (Table 1). The SelAC species tree performed poorly relative to FMutSel0 (Fig. S2.3.3; Tables 2, S2.3.2), despite its incorporation of GTR nested within a codon and protein model (Beaulieu, et al. 2019). Perhaps this implies that aspects of the model require improvement, at least when applying it to individual genes. SelAC is designed to estimate parameters from whole multi-locus datasets, so limiting its optimization to single genes is a potential limitation to the current implementation of the model. We should also note that recent updates to the SelAC package (which also implements FMutSel0 and other models) have accelerated its computation speed (O’Meara, unpub. data) allowing improved results via more thorough optimizations.

There are some other caveats to highlight. First, our original hypothesis favored SelAC because part of our research group developed this model and found it to outperform others (including FMutSel0; Beaulieu, et al. 2019). This is contrary to our results and possible reasons why are discussed above. Next, loci were chosen that had minimal rate heterogeneity to minimize modelling issues (Yang 1996; Chira and Thomas 2016), but there are other factors to consider when choosing optimal loci (Meyer, et al. 2011; Misof, et al. 2013; Doyle, et al. 2015; Steel and Leuenberger 2017; Evangelista, et al. 2019 unpublished data). Minimizing among site rate heterogeneity could be disadvantageous if it resulted in higher rate heterogeneity among lineages (Zhong, et al. 2011). This may be an additional reason why FMutSel0 was found to be superior to SelAC because the latter model would be expected to outperform FMutSel0 when data is more heterogeneous due to its use of multiple substitution matrices (pers. obs. M.A. Gilchrist). However, we found no difference (Z-test, p>0.05) in the performance of either FMutSel0 or SelAC in identifying the least discordant gene tree among the 14 least and 14 most heterogeneous loci (Table S2.1.2). Finally, based on our examination of gene trees, it is our opinion that many of the errors would be classified as long branch effects (Wägele and Mayer 2007; Roch, et al. in press) and thus merit a similar but more nuanced interpretation of the results. Given the difficulty in objectively identifying long branch effects in a tree (e.g., Kück, et al. 2012; Evangelista, et al. 2018), let alone a set of gene trees, we do not attempt at quantifying it.

### Implications for Blaberidae

These findings have some novel implications for the phylogeny of Blaberidae, but interpreting them requires some context. Evangelista, et al. (2019 unpublished data) inferred a phylogeny of Blaberoidea, including 48 species of Blaberidae, with a concatenated analysis of 100 genomic loci. That study went to great lengths to infer a robust tree by identifying the most informative genomic loci. However, being a concatenation analysis, it is expected to poorly recover anomalous nodes (Mendes and Hahn 2018). Both that study and this one show that that many nodes within Blaberidae are near the anomaly zone (i.e., the species tree topology is not found in most of the gene trees; low rates of independent gene tree validation and internode certainty).Yet, the species trees recovered here also require cautious interpretations as they only leverage 40 genomic loci.

Comparison with the baseline trees provides evidence for specific relationships being attributed to congruent or discordant (i.e., possibly erroneous) gene tree topologies. Relationships recovered in the test species trees that are possibly due to gene tree error are: Asian-Epilamprinae being sister to (Pycnoscelinae + Asian-Perisphaerinae); *Laxta* being sister to Panesthiinae; and Paranauphoetinae being sister to Oxyhaloinae. However, none of these were recovered in the most plausible tree (FMutSel0; Fig. 6). Conversely, minimizing discordance results in the recovery of two other deep relationships: *Paraplecta minutissima* as sister to Oxyhaloinae; and Diplopterinae as sister to (*Paraplecta minutissima* + Oxyhaloinae). These were only found in Est.ST.1 and the FmutSel0 species tree. Encouragingly, these relationships have a strong morphological argument to support them (supplementary material S2.3), one was seen in prior studies (Legendre, et al. 2017; Bourguignon, et al. 2018; Evangelista, et al. 2019), and they were recovered in the FMutSel0 species tree with 0.79 local posterior probability quadripartition support. As discussed in supplementary material S2.3, this topology suggests the ancestor of this clade had a coleopteroid gestalt.

**Figure 6.**
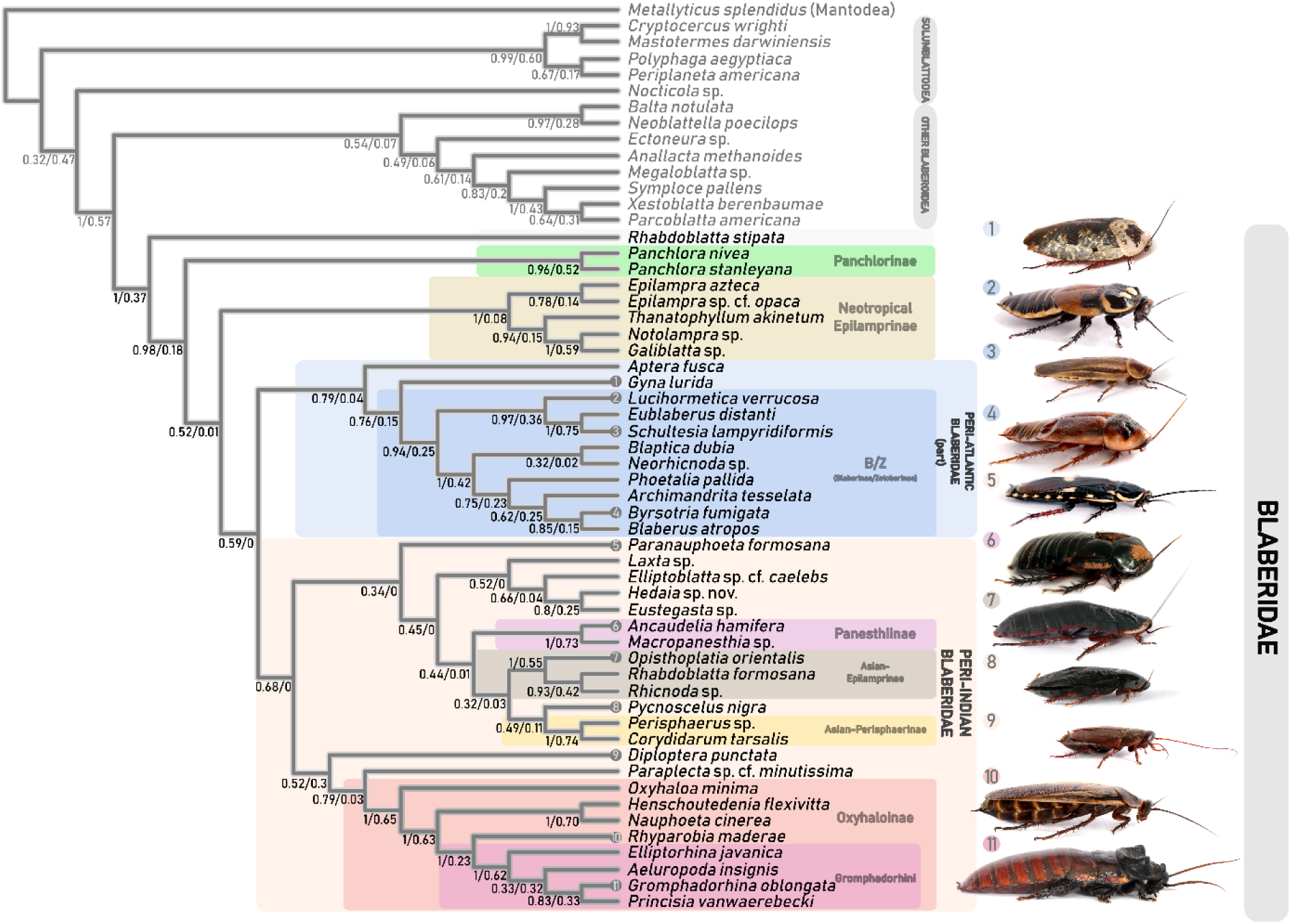
Species tree of Blaberidae. Inferred from an ASTRAL analysis of gene trees of 40 loci as chosen by the FMutSel0 model. Values on edges are local quadripartition posterior probability / predictive power of the bipartition. The second value is calculated as the frequency each bipartition occurs in a set of 120 gene trees inferred from 60 independent loci under two models (GTR, ECM). Numbered circles correspond tips to photographs (exceptions: 1 - *Gyna caffrorum;* 2 - *Diploptera minor*).

Some relationships are consistently recovered in coalescent analyses but are never found with concatenation - suggesting that GTST discordance needs to be accounted for on these nodes. This was the case in finding *Aptera fusca* as sister to *Gyna lurida* + B/Z (Blaberinae/Zetoborinae; pp = 0.79); and *Princisia vanwaerebecki* sister to *Gromphadorhina oblongata* (pp = 0.83). The latter is a relationship suspected by morphological similarity (pers. comm. George Beccaloni), as opposed to the relationship proposed by concatenation (Evangelista, et al. 2019 unpublished data).

The FMutSel0 tree (Fig. 6), which was the most plausible species tree, has Panchlorinae as sister to the remaining Blaberidae except *Rhabdoblatta stipata* (pp = 0.52). This is not congruent with other phylogenomic studies [i.e., Panchlorinae within Peri-Atlantic Blaberidae (Bourguignon, et al. 2018; Evangelista, et al. 2019; Evangelista, et al. 2019 unpublished data)] but has been seen before (e.g., Legendre, et al. 2015; Evangelista, et al. 2018). It is interesting that phylogenomic concatenation studies recover Panchlorinae within Peri-Atlantic Blaberidae since this relationship was absent in all 120 independent gene trees used to evaluate the species tree (Table S2.3.2). This appears to be a case for the emergent/hidden support phenomenon of concatenation (Gatesy and Springer 2014). The relationship recovered here is fairly common among gene trees (occurring 17.5% of the time) but was not in any other species tree. A more common gene tree topology (29.3%) was Panchlorinae as sister to *Diploptera punctata.* This was recovered in previous studies (reviewed in Evangelista, et al. 2018) but may be a long-branch artifact. This topology is found in the SelAC, Est.ST.2, and max. discordance species trees.

Evangelista, et al. (2019), with small taxon sampling, showed that the major lineages of extant Blaberidae (*Panchlora, Gyna lurida*, B/Z, *Diploptera punctata*, Oxyhaloinae) originated within a span of ∼35 Myr. after the origin of crown-Blaberidae. Our tree (Fig. 6) suggests two reasons for potentially revising this perception of Blaberidae’s diversification. First, we show that between four and eight additional speciation events occurring along the backbone of Blaberidae during this diversification period (diversification of Peri-Indian Blaberidae; the splitting of *Paraplecta minutissima* and *Oxyhaloa minima* from other Oxyhaloinae; splitting of neotropical-Epilamprinae and *Aptera fusca* from their respective sister groups). This could drive a node density effect (Venditti, et al. 2006; Hugall and Lee 2007) and increase the time since divergence and unpredictably change the duration of the clade’s diversification. Second, Evangelista, et al. (2019) did not include *Rhabdoblatta stipata* or any other African-Epilamprinae, which Evangelista, et al. (2019 unpublished data) and our study found to be sister to the remaining Blaberidae with strong support (pp = 1). Including these would have shifted the location of the fossil calibration relative to other Blaberidae lineages and thus may have narrowed the timeframe of diversification. A novel diversification analysis is needed to clarify these issues.

### Conclusions

In summary, we first found that gene trees estimated with the GTR model in IQ-TREE tend to be less discordant with the concatenation tree compared to those estimated with the ECM codon model. Ranking the gene trees by their likelihood under two selection-based codon models, we found that discordance is probably overestimated by any given simple model, but discordance is never absent. The species tree inferred from gene trees chosen by FMutSel0 were determined to be more internally consistent, more predictive of independent data, and more congruent with morphological hypotheses. This supports the conclusions of the FMutSel0 model: that gene tree species tree discordance is overestimated by other models. The amount of discord in the gene tree sample has a biologically meaningful effect on species tree topology. We identify eight deep lineages in our tree are specifically affected by discord among gene trees.

## Acknowledgements

Great thanks to Sabrina Simon, Benjamin Wipfler, Karen Meusemann and the 1KITE consortium for providing early access to datasets. Thank you to Luna Sanchez Reyes, Orlando Schwery, David Babst, Sam Borstein, Cedric Landerer and Jeremy Beaulieu for providing critical feedback and assistance. This study was funded by NSF award no.’s 1608559 and 1355033.

## Author contributions

DAE conceived the study and developed the workflow with guidance from MG and BO. FL and DAE determined taxon sampling. MG and BO conceived the statistical tests and tree search regimes. DAE composed the figures and wrote the manuscript with assistance from MG, BO, and FL. DAE photographed specimens.

