## Supplementary Material for "The Implications of Over-Estimating Gene Tree Discordance on a Rapid-Radiation Species Tree (Blattodea: Blaberidae)"

Submitted to BioRxiv 28 - July- 2019 by

Dominic A. Evangelista, Michael Gilchrist, Frederic Legendre, Brian O'Meara

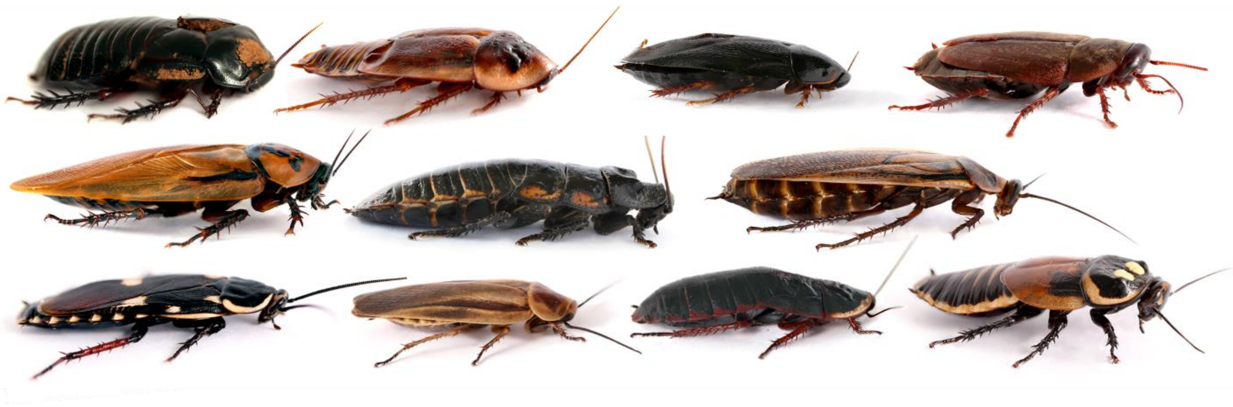

### S.1 Justification of methods

#### S.1.1 Choice of evolutionary models

The goal of inferring gene trees under different models is to examine cases where there are multiple plausible topologies for a given locus and come to conclusions about differing discordance with the species tree. For this comparison to be biologically meaningful both alternatives must be robust hypotheses given different assumptions. We chose the first hypothesis to be dictated by the assumptions of the GTR model. GTR is an extremely widely used nucleotide model, as evidenced by the popularity of phylogenetic software that only implements this model (Stamatakis 2014). It is the most complex and versatile of nucleotide models (Tavaré and Miura 1986) and thus should yield meaningful evolutionary histories in most cases. Additionally, GTR is the only nucleotide model that could be implemented in both our tree inference software (IQ-TREE) and our tree testing software (SelAC). We decided the second hypothesis should also be meaningful but should incorporate codon-level information, since codon models are thought to be superior to nucleotide models (e.g., Goldman and Yang 1994; Wang et al. 2014; Arenas 2015; Doud et al. 2015; Sealfon et al. 2015). However, there is little consensus on which codon model would be most appropriate. In order to ensure that the codon model was meaningful, we allowed some flexibility. Before each inference of the second hypothesis IQ-TREE's built in model-finder determined if ECMS05 (Schneider et al. 2005) or ECMK07 (Kosiol et al. 2007) and which frequency model was most appropriate. These models differ in that: GTR only considers nucleotide changes and estimates substitution frequencies from the data (Tavaré and Miura 1986), both ECM models consider codon structure, ECMS05 uses observed substitution rates from a large vertebrate dataset (Schneider et al. 2005), and ECMK07 uses physicochemical properties of amino-acids to estimate codon evolutionary rates (Kosiol et al. 2007). ECMK07 also allows for instantaneous doublet or triplet mutations to occur (Kosiol et al. 2007). Surprisingly, ECMS05 was chosen as superior for all loci.

The goal of testing alternative gene tree topologies with SelAC, and FMutSel0 were to use the most biologically plausibly models, despite their complexity. The incredible complexity of protein evolution justifies such approaches considering that many parameters varying across individual sites. For instance, evolutionary rate of any given amino acid is known to vary, increasing if it is: on the protein surface, contributing to protein flexibility, in a region of protein structural disorder, in a lowly expressed protein, not necessary to maintain a stable structure, or not playing a role in the active region of the protein (Echave et al. 2016). Determining evolutionary dynamics based on these mechanistic predictors requires biophysical models (Echave et al. 2016), which SelAC does not explicitly take into consideration but can approximate through heterogeneity of the strength of selection. Related to variable rate evolution of amino acid sites, there is also evolutionary preference for certain amino-acid states at specific sites (Wang et al. 2014; Doud et al. 2015; Risso et al. 2015). It has also been hypothesized that sites with changing amino-acid preferences positionally correlate with sites that are fast evolving but this has not always been demonstrated (Doud et al. 2015). Both SelAC and FMutSel0 consider site specific amino-acid preferences, which tend to strongly outperform models lacking such parameters (Doud et al. 2015). Finally, both rate of evolution and the site-specific preferences for certain amino-acids have epistatic effects (Hoehn et al. 2017) and these features may or may not change over time (Risso et al. 2015; Usmanova et al. 2015). Again, SelAC does not model these explicitly, but provides a framework under which the resulting patterns can be modelled.

#### S.1.2 Control nodes and justification

One method by which we can assess the plausibility for a species tree is to judge whether it contains nodes, or relationships that have been well-established with strong support (i.e., Shen et al. 2017). In other words, an estimated species tree should reliably recover uncontroversial relationships. Below are eight relationships among the taxa we included in our analyses that we deem uncontroversial. The prior evidence to support this is discussed.

### S.2 Supplemental results

#### S2.1 Model tests: 66 loci

We did preliminary tree optimizations for 66 loci. They had on average, 45 taxa (min. 34, max. 51) per locus. Branch lengths for each of the three trees for each locus were optimized again using the SelAC software package (Beaulieu et al. 2019). We fit: GTR+G4+FO, SelAC+GTR+G4+FO+amino acid optimization (AAO), and MutSel+GTR+G4+FO+AAO. In both cases, estimated branch lengths from IQ-TREE were used as starting values and optimization chain was run for a single set of initial conditions with six iterations of 1000 evaluations each and the criteria max.tol.edges = 1.4 and tol.step = 2.3, parallelized over two processors. The log-likelihood (lnL) was used to determine which gene tree was preferred by which model.

The results from testing all 66 loci with three models in the SelAC software package are as follows. The simplest evolutionary model (GTR) assigned the highest lnLikelihood (lnL) to estimated topology 1 70% of the time. The SelAC and FMutSel0 models also assigned the highest lnL to estimated topology 1 a majority of the time (59-64% of the time). Each model assigned the highest lnL to a less discordant topology more often than to a more discordant topology but only FMutSel0 did so in a manner that was statistically significant compared to random tree choice (i.e., $p<0.05$). Despite the overall trend towards reducing gene-tree discordance, the species-tree topology was rejected in an overwhelming majority of tests. The statistical significance of model family on a topology's optimality under the three models was more apparent. GTR was much more likely ($p<0.05$) to choose estimated topology 1 than estimated topology 2 (presumably because estimated topology 1 was inferred with GTR). FMutSel0 ($P<0.05$) was less likely to choose estimated topology 2 (estimated with another codon model). SelAC was indistinguishable from randomness when comparing against either model family.

##### *Table S2.1.1*

The best trees chosen for all loci tested. Statistical significance is denoted the an * and was determined using a Z-Test with alpha = 0.05. FMutSel0 significantly chose less discordant gene-trees more often than more discordant ones. However, FMutSel0 and GTR both showed a significant bias for trees generated under a certain model family (i.e., they preferred estimated topology 1 over estimated topology 2).

| | n | Best LnL: sp. tree top. | Best LnL: Est. top. 1 | Best LnL: Est. top. 2 | Proportion of best: Concat. tree topology | Proportion of best: Same model family | Proportion of best: Less discord |
|---|---|---|---|---|---|---|---|
| **SelAC** | 66 | 0 | 39 | 27 | 0.00 | 0.41 | 0.61 |
| **FMutSel0** | 64 | 0 | 41 | 23 | 0.00 | 0.36 * | 0.66 * |
| **GTR** | 67 | 1 | 46 | 20 | 0.02 | 0.70 * | 0.62 |

***Table S2.1.2***

Comparison of gene tree optimizations of loci with different levels of among site rate heterogeneity within 66 loci. Rate heterogeneity is calculated by determining the mean number of rate categories per nucleotide [specifics of calculation in Evangelista et al. 2019 unpublished data]. The values in the table show how often the models correctly identified the least discordant tree. P-values show the probability that the least discordant tree was identified at the same rate in both heterogeneity sets as determined by a Z-Test.

| | % of hits on the least discordant gene tree | | |
|---|---|---|---|
| | **Low Heterogeneity** | **High Heterogeneity** | ***p*** |
| **SelAC** | 43% | 53% | 0.87 |
| **FMutSel0** | 48% | 57% | 0.28 |
| **GTR** | 48% | 38% | 0.09 |
| *n* | *14* | *14* | |

#### S2.2 Comparison of six species trees

##### *Figure S2.2.1*

The full concatenation tree topology compared to six species-trees inferred in this study. (a) The concatenation species-tree topology inferred in Evangelista et al. (2019 unpublished data) with taxa trimmed. (b-g) "Cophylo" plots of six species-trees against the concatenation tree shown in (a). (b,c) Species-trees inferred from all the most congruent 40 gene-trees and the most incongruent 40 gene-trees respectively. (d, e) Species-trees inferred from all 40 gene-trees of Est.Top.1 and Est.Top.2 respectively. (f, g) Species-trees inferred from all 40 gene-trees selected by FMutSel0 and SelAC respectively. Blue pies on nodes show local posterior probability support. (h) Symmetrical matrix of Robinson-Foulds (RF) distances among all species trees. Colors emphasize high (red) and low (blue) RF distances.

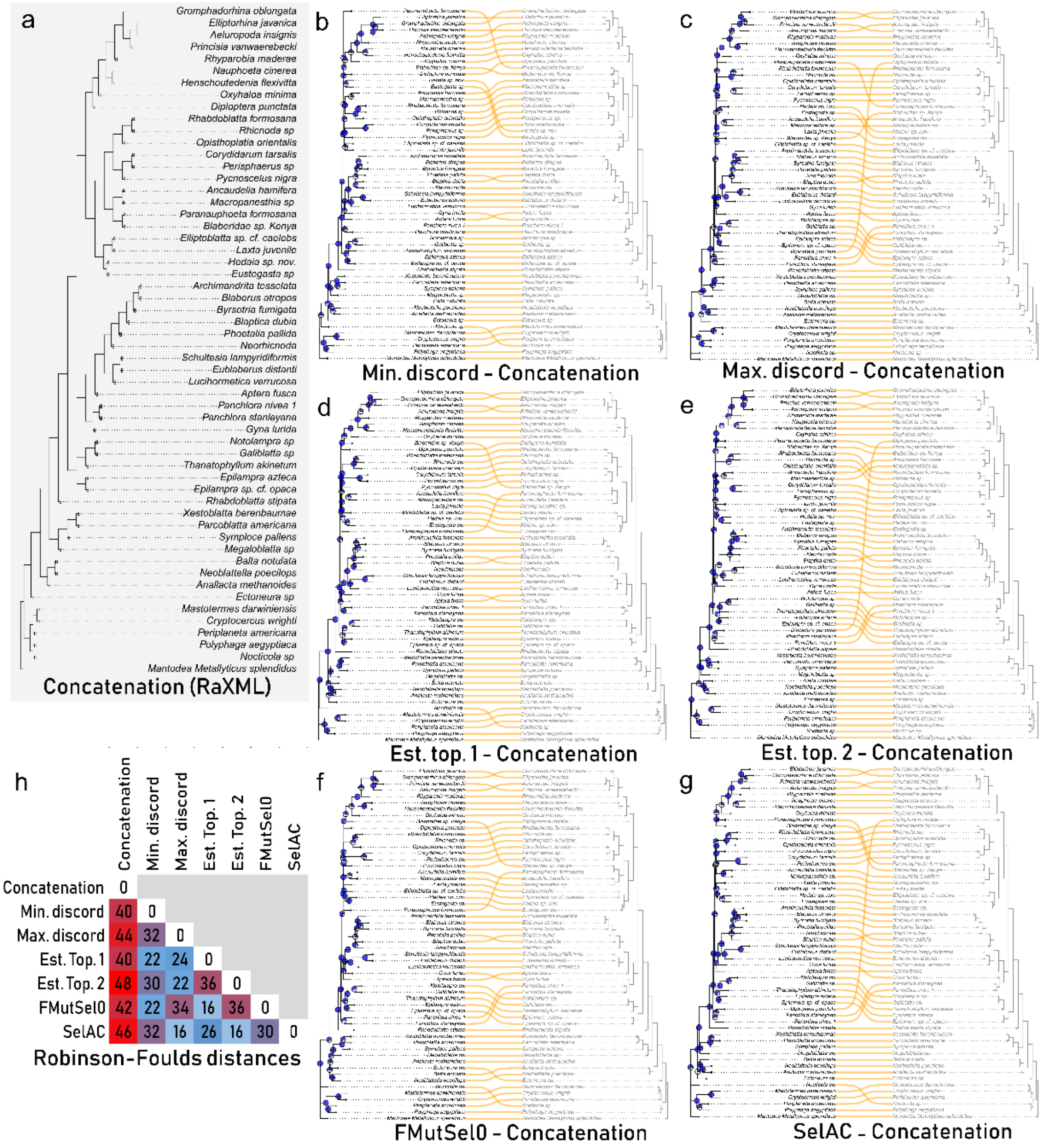

#### S2.3 Morphological and other support for recovered relationships

One relationship recovered in both the concatenation tree and most of the coalescent trees was Pycnoscelinae as sister to Asian Perisphaerinae. While the local posterior probability of the relationship is low (0.49), this relationship appears in independent gene trees at a high rate (10.8%) and is also congruent with the concatenation topology. The Est.Top.2 and SelAC species trees had different relationships. Roth (1973a) showed morphological evidence for a relationship between Pycnoscelinae, Diplopterinae and Oxyhaloinae (but see McKittrick 1964) but we never found this topology. A relationship with Asian Perisphaerinae or Diplopterinae both make sense biogeographically − all are distributed in South-East Asia. There has previously been no consistent molecular (Djernæs et al. 2012; Legendre et al. 2014; Legendre et al. 2015; Legendre et al. 2017; Bourguignon et al. 2018; Evangelista et al. 2018) hypothesis for Pycnoscelinae.

Some relationships were consistent across the coalescent trees but differed with the concatenation trees (Figs. S2.2.1, S2.3.3). These are prime suspects for errors in the concatenation species-tree due to ILS. The differences between the coalescent species-tree and the concatenation species tree are in four locations.

First, the deep relationships among Peri-Atlantic Blaberidae. All the coalescent trees agreed that Gyninae (only *Gyna lurida* was sampled) was sister to the Blaberidae/Zetoborinae (BZ) complex and *Aptera fusca* was sister to both of these and finally Panchlorinae is sister to the remaining Peri-Atlantic Blaberidae. In contrast, our concatenation tree, and that of a recent phylotranscriptomic study (Evangelista et al. 2019), showed Gyninae as more closely related to Panchlorinae [although Evangelista et al. (2019) did not sample *Aptera fusca*]. Little morphological systematic work has been done to clarify the positions of these taxa (but see Grandcolas 1993). Both topologies offer equally parsimonious biogeographical scenarios (two transitions each) but depend on the timing of the splits in coordination with the drift of S. America away from Africa. However, Legendre et al. (2017) and unpublished data (FL) indicate that more taxa may be needed to precisely clarify this region of the tree (e.g., *Gynopeltis* spp., other putative Gyninae, and African Perisphaerinae).

Second, ASTRAL-III always recovered *Blaberus atropos* as sister to *Byrsotria fumigata* instead of sister to *Archimandrita tessellata,* which is the concatenation hypothesis. Roth (1970c) indicates that *Blaberus* and *Archimandrita* are most closely related (also see Legendre et al. 2017). Thus, the ASTRAL hypotheses are at odds with the previous hypotheses for these genera.

Third is the position of *Phoetalia pallida* and *Blaptica dubia*, which was sister to *Byrsotria fumigata + Archimandrita tessellata + Blaberus atropos* in all the coalescent trees while it was sister to *Blaptica dubia + Byrsotria fumigata + Blaberus atropos + Archimandrita tessellata* in the concatenation tree. The position of *Phoetalia* was previously debated by McKittrick (1964), Roth (1970c) and Legendre et al. (2017) but the debate was largely concerning its relationship to the “Blaberinae” complex or the “Zetoborinae” and not its position relative to *Byrsotria*.

Finally, *Gromphadorhina oblongata* was sister to *Princisia vanwaerebecki* in all ASTRAL trees but sister to *Aeluropoda insignis + Elliptorhina javanica + Princisia vanwaerebecki* in the concatenation tree. *Princisia* and *Gromphadorhina* are very morphologically similar (two robust pronotal horns and a raised anterior pronotal margin) and thus are thought to be closely related (pers. obs. Evangelista; G. Beccaloni pers. comm.; also see Legendre et al. 2017). The putative synapomorphies uniting *Princisia* and *Gromphadorhina* could be symplesiomorhies though. Particularly considering the earlier branching of *Leozehntnera* (Legendre et al. 2017), which has a similar pronotal morphology. Genital morphology is relatively conserved in this group, and are thus mostly uninformative.

Relationships recovered in the positive control (congruence) coalescent tree but not in the negative control tree (incongruence) are in line with our hypothesis that most gene tree discordance is due to gene tree error and not ILS (i.e., the least discordant gene tree is probably the correct gene tree). These cases are only seen in the Est.Top.1 species tree and the FMutSel0 gene tree, with respect to the closest relatives of Oxyhaloinae. The congruence, Est.Top.1, and FMutSel0 species trees have *Paraplecta minutissima* as sister to Oxyhaloinae. *Paraplecta* was suggested to be more closely related to Perisphaeriinae based on the excavation of the right side of the subgenital plate (Roth 1995; also see Roth 1973b). However, the excavation is hooked like in Oxyhaloinae (who have hooks on both the left and right sides). The same trees have Diplopterinae (*Diploptera punctata*) as sister to *Paraplecta minutissima* + Oxyhaloinae. The alternative (seen in the Est.Top.2 and SelAC trees) is Diplopterinae as sister to Panchlorinae. This alternative was also found in some other molecule-based studies (Legendre et al. 2017; Evangelista et al. 2018) but this may be due to long-branch attraction. A close relationship between Diplopterinae and Oxyhaloinae was a prevailing morphological hypothesis in precladistic morphological studies (reviewed in Li and Wang 2015) and was also found in recent phylogenomic studies (Bourguignon et al. 2018; Evangelista et al. 2019; Evangelista et al. 2019 unpublished data). Since we always recovered *Oxyhaloa duesta* as sister to the remaining Oxyhaloinae, our tree suggests that a "coleopteriod" body form (dorso-ventrally thickened, strongly tegmenized or elytriform forewings, greatest body width more than half medial body length; seen in *Diploptera punctata*, *Paraplecta minutissima*, and *Oxyhaloa duesta*) is the plesiomorphic state to Oxyhaloinae and a the more traditionally roachoid formed (dorso-ventrally flattened; moderately or lightly tegminized forewings; greatest body width less than half medial body length) is autapomorphic in the clade.

Relationships recovered in the negative control (incongruence) coalescent topology and not the positive control (congruence) topology are not in line with our hypothesis. If true, they would indicate that most gene-tree discordance with the species-tree are due to ILS. We see this case first between the incongruence and Est.Top.1 tree with Panesthiinae and *Laxta* sp. and second between the incongruence, Est.Top.2 and SelAC trees with respect to Paranauphoetinae and Oxyhaloinae. In incongruence and Est.Top.1, Panesthiinae and *Laxta* sp. are sister taxa. This is at odds with the monophyly of Perisphaerinae, which is has a morphological basis (Grandcolas 1997; Anisyutkin 2003). However, it could be the case that *Laxta* sp. is correctly positioned as sister to Panesthiinae and the other Perisphaerinae are the ones misplaced (Anisyutkin 2003). *Laxta* has previously been considered as an Epilamprinae by both morphological (Roth 1992) and molecular (Bourguignon et al. 2018) determination. The incongruence, Est.Top.2, and SelAC trees all recovered Paranauphoetinae as sister to Oxyhaloinae. Anisyutkin (2003) proposed that Paranauphoetinae was sister to Perisphaerinae + Panesthiinae with three morphological character states supporting the monophyly of all three taxa and three morphological characters separating Paranauphoetinae from the other two.

We also see that both the concatenation topology and incongruence topology support Asian-Epilamprinae as sister to Pycnoscelinae + Asian-Perisphaerinae. This relationship could be found in both trees due to a common source of error (i.e., signal erosion) in the individual gene alignments and concatenated alignments. It could also be a correct relationship and would thereby support that our hypothesis (most GTST discord is due to error) is incorrect. We discuss above that Pycnoscelinae as sister to Asian-Perisphaerinae was recovered in most trees. However, the only known phenotypic support for this relationship is their shared biogeographical range. The same is true for Asian-Epilamprinae − the taxa we included all reside in S.E. Asia. Given that Epilamprinae (sensu Roth 2003) are prominently distributed in S. America and Africa one might think that the recovered topology represents a conflict with the morphological hypothesis for the group. However, it is already suspected that Epilamprinae is polyphyletic (Legendre et al. 2017; Bourguignon et al. 2018). The lineages in question were only included in one prior molecule-based study where the topology was Asian-Epilamprinae as sister to a clade containing variety of other Blaberidae but with very low support (Legendre et al. 2017). Thus we consider prior studies to be agnostic towards any specific relationship of Asian-Epilamprinae.

***Table S2.3.1***

Plausibility of species trees as determined by approximately unbiased (AU) tests and node support. Each AU test was done with the GTR+G model and a different concatenated dataset: 265 loci without partitioning, 265 loci with codon positions in optimized partitions, and 40 loci with no partitioning. The difference from the best likelihood (ΔlnL), and the p-value (p) is given for each tree in each test. Trees with p<0.005 are considered implausible given the alignment. Plausible test results are marked with *. ASTRAL inferred tree from FMutSel0 gene trees was plausible given a concatenated alignment of 265 loci without partitioning, but all other alignments and trees tested were deemed implausible.

| | | **AU Test Results** | | | | | |
|---|---|---|---|---|---|---|---|
| | | **265 loci, no partitioning** | | **265 loci, codon partitioning** | | **40 loci, no partitioning** | |
| **Species Tree** | **Type** | ΔlnL | p | ΔlnL | p | ΔlnL | p |
| Concatenation | baseline (RaXML) | 0.00 | 1.00* | 0.00 | 1.00 | 0.00 | 0.99 |
| Min. discordance | + control (ASTRAL) | 818.28 | 0.00 | 833.58 | 0.00 | 166.29 | 0.00 |
| Max. discordance | - control (ASTRAL) | 798.65 | 0.00 | 803.62 | 0.00 | 209.72 | 0.00 |
| Est. top. 1 | ASTRAL | 705.93 | 0.00 | 722.85 | 0.00 | 162.70 | 0.00 |
| Est. top. 2 | ASTRAL | 1646.66 | 0.00 | 1660.09 | 0.00 | 318.76 | 0.00 |
| SelAC | ASTRAL | 1576.47 | 0.00 | 1586.14 | 0.00 | 278.33 | 0.00 |
| FMutSel0 | ASTRAL | 833.26 | 0.71* | 862.28 | 0.00 | 96.06 | 0.01 |

***Table S2.3.2***

Frequency of relationships found in 120 independent gene trees. Percentages give the frequency that the defined relationships were recovered in a data set of gene trees inferred from 60 loci not included in the species tree inference. Gene trees were inferred in IQTREE with the two methods described in the main text. Cells are colored by percentage so high percentages are red and low percentages are blue.

| Taxon | Sister to | Frequency in 120 independent gene trees: Est.Top.1 | Est.Top.2 | SelAC | FMutSel0 | Other |
|---|---|---|---|---|---|---|
| *Laxta* | *Elliptoblatta* | | | | | 4.5% |
| | *Elliptoblatta + Hedaia + Eustegasta* | | 0.0% | 0.0% | 0.0% | |
| | Panesthiinae | 3.3% | | | | |
| *Paranuphoeta* | Remaining Peri-Indian Blaberidae | | | | | 0.0% |
| | Remaining Peri-Indian Blaberidae - (Oxyahloinae + *Paraplecta + Diploptera)* | 0.0% | | | | |
| | *Laxta + Elliptoblatta + Hedaia + Eustegasta* + Panesthiinae + Asian-Epilamprinae + *Pycnoscelus* + Asian-Perisphaerinae | | | | 0.0% | |
| | Oxyhaloinae | | 3.3% | 3.3% | | |
| Panesthiinae | Asian-Epilamprinae | | | | | 2.5% |
| | Asian-Perisphaerinae | | 1.7% | | | |
| | Asian-Perisphaerinae + *Pycnoscelus* + Asian-Epilamprinae | | | 0.8% | 0.8% | |
| | *Laxta* | 3.3% | | | | |
| *Elliptoblatta* | *Paraplecta* | | | | | 3.4% |
| | *Laxta* | | | | | 4.5% |
| | *Hedaia + Eustegasta* | 3.6% | 3.6% | 3.6% | 3.6% | |
| *Paraplecta* | Oxyhaloinae | 3.0% | | | 3.0% | |
| | *Paranauphoeta* + Oxyhaloinae | | 0.0% | 0.0% | | |
| Oxyhaloinae | *Paraplecta* | 3.0% | | | 3.0% | |
| | *Paranauphoeta* | | 3.3% | 3.3% | | |
| *Diploptera* | *(Paraplecta + Oxyhaloinae)* | 2.5% | | | 2.5% | |
| | *P. stanleyana* | | 0.0% | 0.0% | | |
| (Paraplecta + Oxyhaloinae) | *Diploptera* | 2.5% | | | 2.5% | |
| Asian-Epilamprinae | *(Pycnoscelus + Perisphaerus + Corydidarum)* | 2.5% | | | 2.5% | |
| | *Pycnoscelus* | | | 1.7% | | |
| | *Paraplecta + Paranauphoeta + Oxyhaloinae* | | 0.0% | | | |
| (Pycnoscelus + Perisphaerus + Corydidarum) | Asian-Epilamprinae | 2.5% | | | 2.5% | |
| *Pycnoscelus* | Asian-Perisphaerinae | 10.8% | | | 10.8% | |
| | Asian-Epilamprinae | | | 1.7% | | |
| | Panesthiinae + Asian-Perisphaerinae | | 0.8% | | | |
| Asian-Perisphaerinae | *Pycnoscelus* | 10.8% | | | 10.8% | |
| | *Pycnoscelus* + Asian-Epilamprinae | | | 2.5% | | |
| | Panesthiinae | | 1.7% | | | |
| Panchlorinae | Sister to all Blaberidae - *R. stipata* | | | | 17.5% | |
| | *Aptera + Gyna* + BZ | 0.0% | | | | |
| | *Diploptera* | | | | | 29.3% |
| *Gyna* | BZ | 15.0% | 15.0% | 15.0% | 15.0% | |
| BZ | *Gyna* | 15.0% | 15.0% | 15.0% | 15.0% | |
| *Aptera* | (*Gyna* + BZ) | 4.0% | 4.0% | 4.0% | 4.0% | |
| *Gyna* + BZ | *Gyna* | 4.0% | 4.0% | 4.0% | 4.0% | |
| *Princisia* | *Gromphadorhina* | 33.3% | 33.3% | 33.3% | 33.3% | |
| *Gromphadorhina* | *Princisia* | 33.3% | 33.3% | 33.3% | 33.3% | |
| *Blaberus* | *Byrsotria* | 15.0% | 15.0% | 15.0% | 15.0% | |
| *Byrsotria* | *Blaberus* | 15.0% | 15.0% | 15.0% | 15.0% | |
| *Phoetalia* | *(Archimandrita + Byrsotria + Blaberus)* | 23.7% | 23.7% | 23.7% | 23.7% | |
| *(Archimandrita + Byrsotria + Blaberus)* | *Phoetalia* | 23.7% | 23.7% | 23.7% | 23.7% | |

***Figure S2.3.3***

Comparison of unique relationships found in each species tree. Each table is organized to differentiate which relationships correspond to a gene-tree population that is highly congruent or highly incongruent, and which result from concatenation or coalescent inference. Each table is organized so that the cells discuss relationships uniquely supported by (from top left to bottom right) only the concatenation tree, the concatenation and congruence tree, only the congruence tree, the concatenation and incongruence tree etc. (a) Explanation of how a relationship could be interpreted. (b) Unique relationships found in the baseline (concatenation) and control trees. (c) The relationships found in test trees. Letters correspond to those in panel (b). (d) Independent evidence for support of each relationship. Each fraction is the number of relationships supported (positive) or discounted (negative) by independent evidence (numerator) and the total number of relationships in that category (denominator). The values separated by semi-colons correspond to: (i) Strong morphological support for relationship; (ii) other support for relationship; (iii) strong morphological support for alternative; (iv) other support for alternative. "Other support" includes support in Evangelista et al. (2019) or other phylogenetic studies (Legendre et al. 2017; Bourguignon et al. 2018; Evangelista et al. 2018; but not Evangelista et al. 2019 unpublished data). Support for most of the relationships is discussed in section S2.3 above.

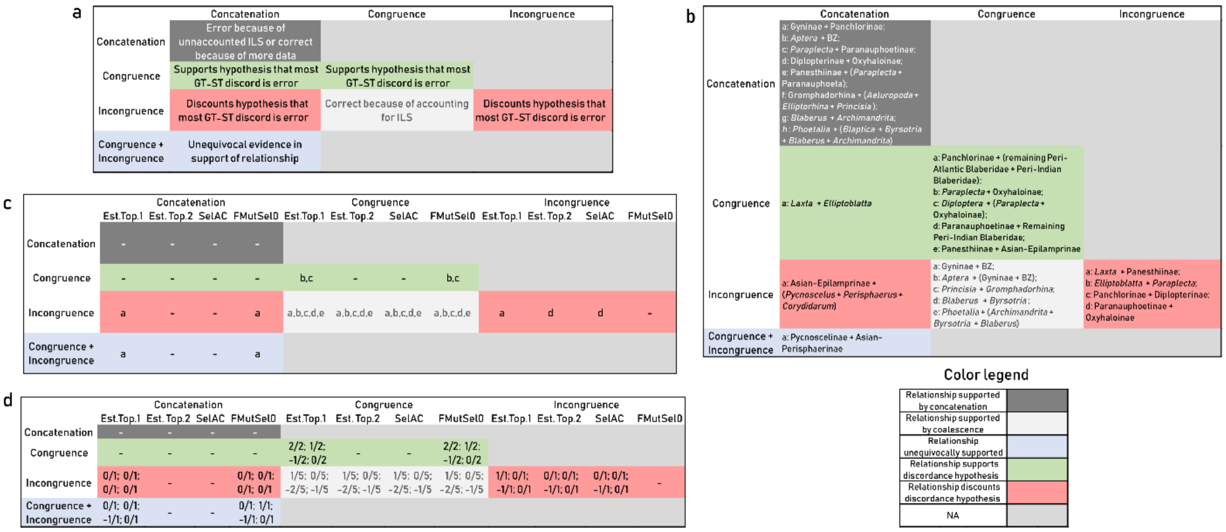
